# A multicellular rosette-mediated collective dendrite extension

**DOI:** 10.1101/303503

**Authors:** Li Fan, Ismar Kovacevic, Maxwell G. Heiman, Zhirong Bao

## Abstract

Coordination of neurite morphogenesis with surrounding tissues is crucial to the establishment of neural circuits, but the underlying cellular and molecular mechanisms remain poorly understood. We show that neurons in a *C. elegans* sensory organ, called the amphid, undergo a collective dendrite extension to initiate formation of the sensory nerve. The amphid neurons first assemble into a multicellular rosette. The vertex of the rosette, which becomes the dendrite tips, is attached to the anteriorly migrating epidermis and carried to the sensory depression, extruding the dendrites away from the neuronal cell bodies. Multiple adhesion molecules including DYF-7, SAX-7, HMR-1 and DLG-1 function redundantly in rosette-to-epidermis attachment. PAR-6 is localized to the rosette vertex and dendrite tips, and promotes DYF-7 localization and dendrite extension. Our results suggest a collective mechanism of neurite extension that is distinct from the classical pioneer-follower model and highlight the role of mechanical cues from surrounding tissues in shaping neurites.

## Highlights

- A multicellular rosette among neurons mediates collective dendrite extension
- Migrating skin carries the rosette vertex to extrude and position dendrites
- Multiple adhesion molecules function redundantly in skin-vertex attachment
- PAR-6 is localized to the vertex and required for attachment and extension

## In brief

Fan *et al.* report a collective dendrite extension mediated by a multicellular rosette, unlike the classical pioneer-follower model. PAR-6 is localized to the rosette vertex and required for the vertex to be carried by migrating skin cells to extrude dendrites.

## INTRODUCTION

Morphogenesis of an organism involves long-range coordination among organs. In the nervous system, sensory neurons may grow long dendrites and integrate the sensory endings to target tissues. The guided outgrowth of neurites needs to be coordinated with other neurons as well as surrounding tissues (Dong, et al., 2015; Lefebvre, et al., 2015; Chao, et al., 2009). In particular, many neurites bundle or fasciculate to form nerves. The pioneer-follower model provides a simple developmental mechanism to explain this organization. The pioneer neuron extends its growth cone first to explore the environment and interact with chemotropic signals and environmental guideposts in order to establish a path to its target. The follower neurons respond to cues from the pioneer (Tamariz and Varela-Echavarria, 2015). Multiple mechanisms coordinate the interaction between the pioneer and the followers, such as selective fasciculation (Hayashi, et al., 2014; Hutter, 2003; Lin, et al., 1994), or juxtaparacrine signals (Jaworski and Tessier-Lavigne, 2012).

The amphids are a bilaterally symmetric pair of sensory organs in the nematode *Caenorhabditis elegans*. Each amphid consists of 12 sensory neurons and two glia cells, namely the sheath and the socket cells. The 12 dendrites in an amphid form a sensory nerve, which extends from the neck of the worm where the cell bodies are situated to the nose tip where most of the ciliated endings of the dendrites are exposed to the environment. An earlier study showed that these dendrites undergo retrograde extension, where the dendritic tips are anchored in place at the embryonic nose known as the sensory depression while the cell bodies move posteriorly, extending the dendrite behind them (Heiman and Shaham, 2009). The anchoring requires DYF-7 and DEX-1, which likely assemble a matrix in the extracellular environment (Heiman and Shaham, 2009).

Here, we report how amphid neurons in *C. elegans* interact with migrating skins to coordinate dendrite extension with organismal morphogenesis, prior to their elongation by retrograde extension. The amphid neurons form a multicellular rosette along with the sheath and socket cells. The vertex of the rosette, which becomes the dendrite tips, is attached to the anteriorly migrating epidermis and carried to the sensory depression, extruding the dendrites away from the neuronal cell bodies. Abolishing epidermis migration by RNAi of *elt-1*, a key transcription factor for epidermal fate, abolishes the extension of the amphid dendrites without affecting rosette formation. DYF-7 is localized at the rosette vertex. In contrast to subsequent retrograde extension which shows complete dependence on DYF-7, loss-of-function phenotypes suggest that DYF-7 acts redundantly with SAX-7/L1CAM, HMR-1/Cadherin and DLG-1/Discs to mediate attachment of the rosette vertex to the migrating epidermis. We further show that PAR-6 is localized to the rosette vertex, and promotes DYF-7 localization, as well as the attachment to the epidermis and dendrite extension. Our study reveals a rosette-mediated mechanism for collective neurite outgrowth and nerve formation, in contrast to the classical pioneer-follower model, and highlights a novel role for neuron-skin interactions in dendrite extension.

## RESULTS

### Amphid neurons form a multicellular rosette before dendrite extension

Using phalloidin staining, we observed a multicellular rosette at the comma stage of *C. elegans* embryogenesis (Fig. 1A). The age of the embryo and the position of the observed rosette suggested that these could be the amphid neurons. To determine the identity of these cells and understand the developmental function of the rosette, we generated a strain using the *cnd-1*/NeuroD promoter driving PH::GFP to label neurons (Shah, et al., 2017). 3D, time lapse imaging and cell lineage tracing with a ubiquitously expressed histone::mCherry (Santella, et al., 2014; Santella, et al., 2010) showed that the rosette is indeed formed by the amphid neurons (Fig. 1B). Specifically, the *cnd-1*p::PH::GFP marker labels 9 of the 12 amphid neurons (ADF, ASE, ASG, ASH, ASI, ASJ, AWA, AWB, AWC). The other 3 amphid neurons are not labeled, but the relative position of their nuclei suggested that they are also part of the rosette. In addition, the sheath and socket cells are labeled and engaged in the rosette. Surprisingly, the marker showed that five other neurons (AIB, AVB, AUA, RIV, URB) are also engaged in the rosette. However, judging by cell shape changes, these non-amphid neurons disengage from the rosette about 40 min later. The significance of this transient engagement is not known.

**Figure 1.**
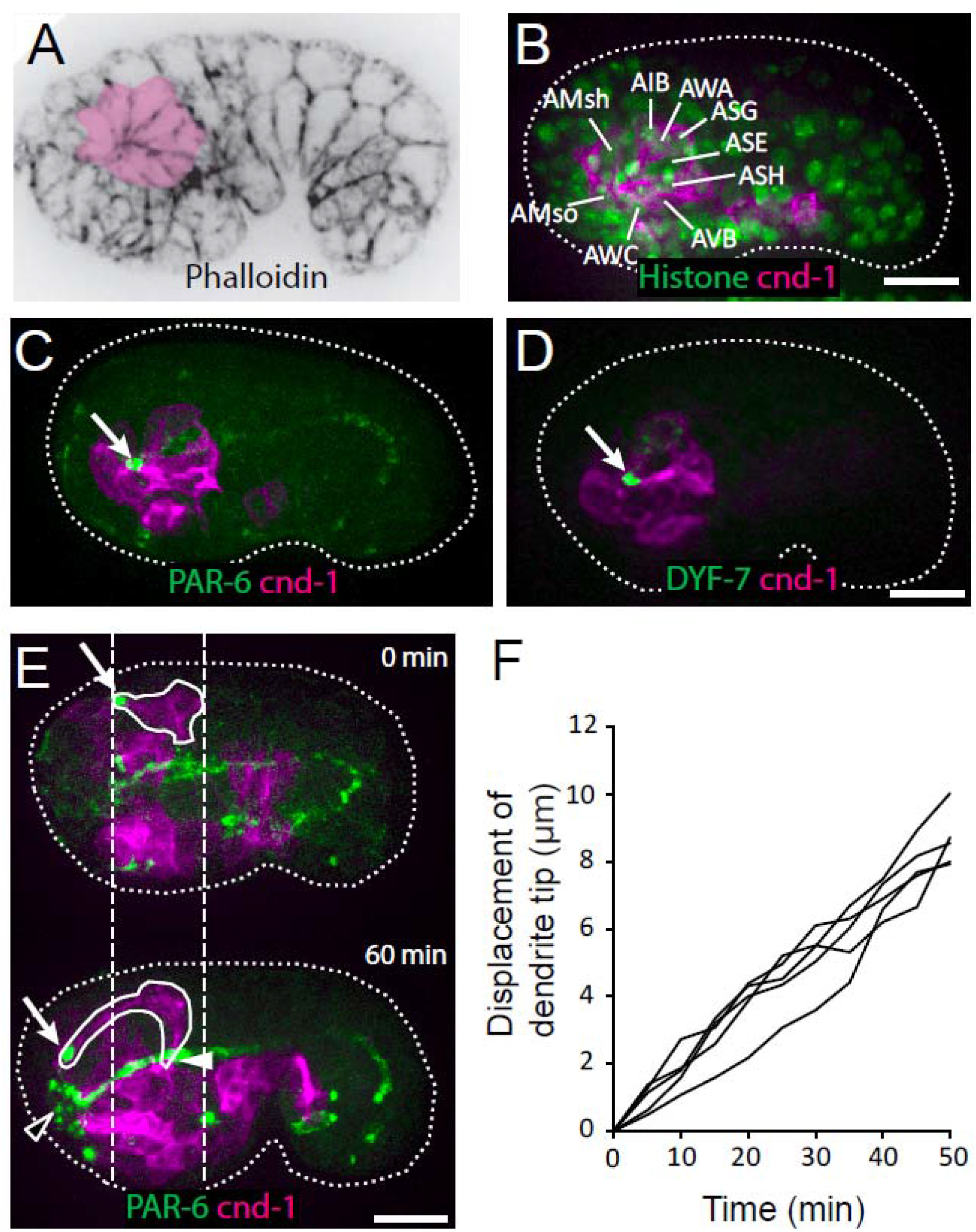
Multicellular rosette precedes collective dendrite extension. (A, B) Amphid neurons form a multicellular rosette. (A) Image of embryo prior to amphid dendrite extension stained with phalloidin. A multicellular rosette is shaded in pink. In this and all subsequent figures, embryos are oriented with anterior to the left. (B) *cnd-1*p::PH::GFP labels amphid neurons in the rosette. Lineage-derived cell identities are shown. (C, D) PAR-6 and DYF-7 are localized to the rosette vertex. *cnd-1*p::PH::mCherry labels neurons. (E) Amphid dendrite tip migrates anteriorly. PAR-6 accumulates in the dendrite tip. *cnd-1*p::PH::mCherry labels neurons. Vertical dashed lines show initial position of amphid dendrite tip and posterior extent of amphid cell bodies. Arrows indicate dendrite tip. Closed arrowhead indicates amphid axon commissure. Open arrowhead indicates sensory depression. (F) Measured displacement of amphid dendrite tips from five embryos during dendrite extension. Scale bars in A-E, 10 μm.

We found that PAR-6 is localized to the center of the rosette (Fig. 1C), suggesting that the rosette is a polarized structure. Furthermore, DYF-7, which is required for anchoring the dendrite tips during retrograde extension (Heiman and Shaham, 2009), is also localized to the rosette center (Fig. 1D).

In the next 60 to 90 minutes, dendrites grow from the rosette center. The dendrite tips migrate anteriorly to the sensory depression. The dendrites stay in a tight bundle, with PAR-6 and DYF-7 localized at their tips (Fig. 1E and F, Movie S1). During this period, the cell bodies remain largely stationary (dashed lines in Fig. 1E). Posterior movement of the cell bodies occurs at later stages (Fig. S1). These results suggest that a phase of anterior extension precedes the previously characterized retrograde extension.

### The rosette vertex is carried by the migrating epidermis

At this developmental stage, the epidermal cells are known to migrate anteriorly to enclose the head (Chisholm and Hardin, 2005). To examine the relationship between the anterior movement of the epidermal cells and the anterior extension of the amphid dendrites, we conducted 3D, time lapse imaging, using an mCherry::PAR-6 reporter (Zonies, et al., 2010) to label the rosette vertex and dendrite tips and a DLG-1::GFP to label the junctions in the epidermal cells (Totong, et al., 2007; Firestein and Rongo, 2001). We found that prior to dendrite extension, the rosette vertex is aligned with the leading edge of the migrating epidermis, specifically the hyp5 cell. As the epidermis migrates anteriorly, an indentation can be observed at the leading edge of hyp5. The dendrite tips are situated in the indentation (Fig. 2A). Furthermore, the anterior movement of the dendrite tips is correlated with the anterior movement of the leading edge of hyp5 (Fig. 2B).

**Figure 2.**
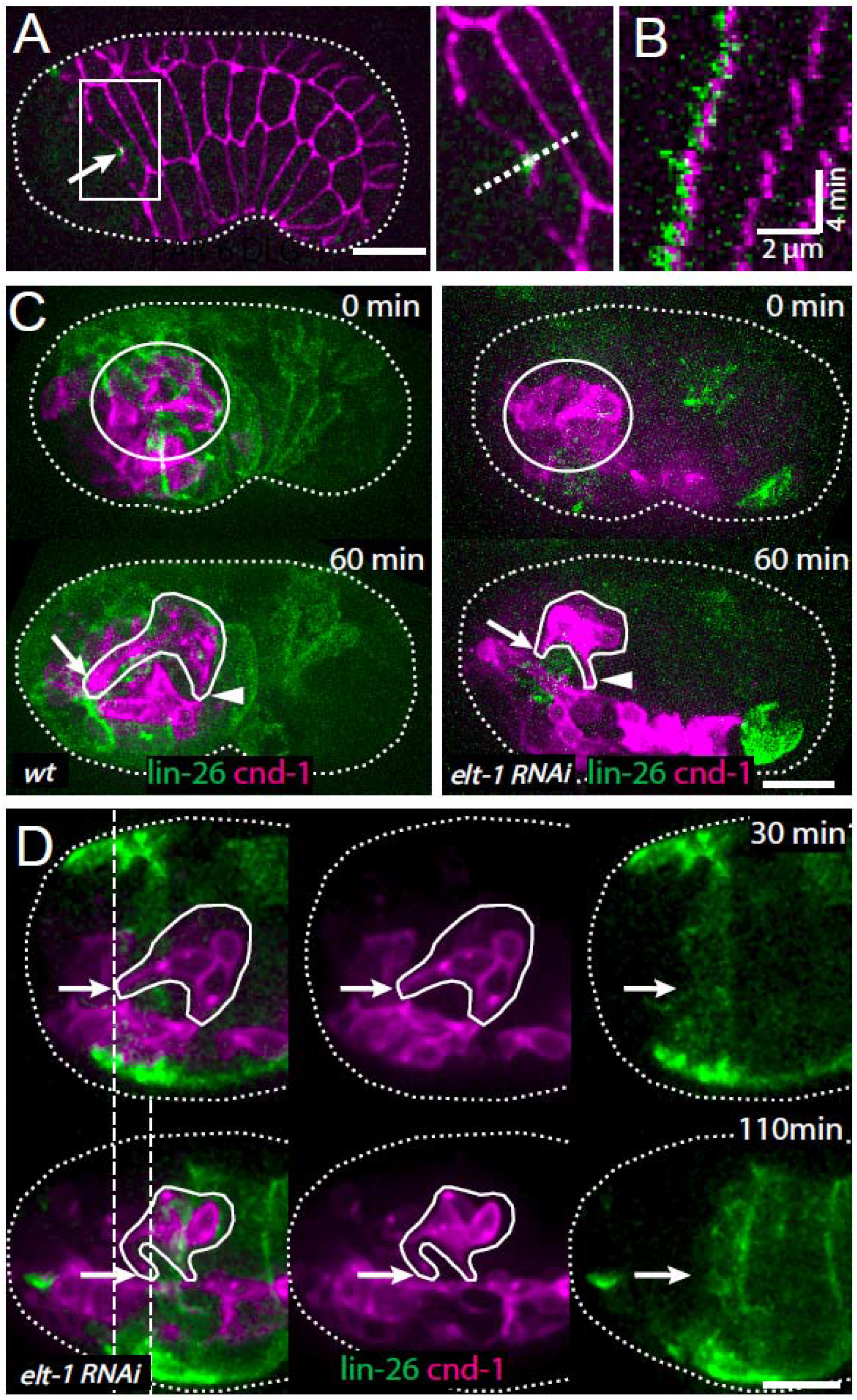
The amphid rosette vertex is attached to the epidermis. (A) PAR-6-labeled amphid tip (arrows) is localized at the hyp5 leading edge. The inset is shown magnified on the right. (B) A kymograph generated along the dashed line in the inset in A, showing the correlated anterior movement of the PAR-6 signal and the leading edge of the epidermal cell. (C) Example of a wild type and an *elt-1(RNAi)* embryo with minimal *lin-26*::GFP expression. In the *elt-1(RNAi)* embryo dendrites failed to extend (arrow), even though the amphid rosette (circle in the upper panel) and the amphid axon commissure both formed normally (arrowheads). (D) A representative *elt-1(RNAi)* embryo with moderate *lin-26*::GFP expression shows a partial anterior migration of the epidermis (upper panel) followed by posterior retraction. Dashed lines mark the leading edge of the epidermis before and after the retraction. The amphid dendrite tips follow the leading edge of the epidermis. Arrows mark the position of the dendrite tip. Scale bars in A-D, 10 μm.

These results suggest that the dendrite tips may be physically attached to the leading edge of hyp5 and carried anteriorly. To test this hypothesis, we sought to perturb the anterior movement of the epidermis. ELT-1 is a GATA transcription factor necessary to specify the epidermal fate (Page, et al., 1997) and activates the expression of its target, *lin-26* (Landmann, et al., 2004). We used *elt-1(RNAi)* to perturb epidermal development, and a *lin-26* promoter driven GFP marker (Gally, et al., 2009) to label the epidermis and assay the consequence. Expression of the *lin-26*::GFP marker was reduced to different levels depending on the efficacy of RNAi. In the embryos with the strongest effect, where minimal *lin-26*::GFP expression remained, dendrite extension was abolished, even though the rosette formed normally (Fig. 2C). In embryos with more moderate RNAi effect, where low levels of *lin-26*::GFP expression were observed, the epidermis made partial anterior movement before retracting posteriorly (dashed lines in Fig. 2D). The dendrite tips remained coupled to the leading edge of the epidermis even as the epidermis retracted posteriorly after failing to enclose the head (Fig. 2D). These results suggest that the dendrite tips are attached to the leading edge of the epidermis and that dendrite extension requires the anterior migration of the epidermis.

### Multiple adhesion molecules function redundantly in amphid dendrite anterior extension

To identify the molecules that mediate the attachment between the dendrites and the epidermis, we examined the localization of several adhesion molecules during dendrite extension. We generated a construct in which the *dyf-7* promoter drives expression of a modified DYF-7 protein that includes a superfolderGFP (sfGFP) tag on its ectodomain, and that completely rescues the dendrite extension defects of a *dyf-7* mutant (data not shown). As previously described, we found that DYF-7 was expressed by amphid neurons and not in epidermis. As expected from its localization to the rosette vertex (Fig. 1B), it localized to the dendrite tips (Fig. 3A). For HMR-1/Cadherin, we used a marker that showed a localization pattern similar to the endogenous pattern as detected by immunostaining and that rescued embryonic lethality of *hmr-1(zu389)* (Achilleos et al., 2010). HMR-1 was observed at the dendrite tips in addition to its known localization at epidermal cell junctions (Fig. 3B). In addition, we examined the localization of SAX-7, a homolog of the vertebrate L1 cell adhesion molecule (L1CAM). SAX-7 has been shown to function redundantly with HMR-1 in blastomere compaction during *C. elegans* gastrulation (Grana, et al., 2010), and to mediate the interaction between the epidermis and the dendrite of the PVD neuron (Dong, et al., 2013; Salzberg, et al., 2013). We used a fosmid-based GFP reporter to examine SAX-7 localization (Diaz-Balzac, et al., 2016). We found that SAX-7 is localized to the epidermal cell junctions and along the amphid dendrites (Fig. 3C).

**Figure 3.**
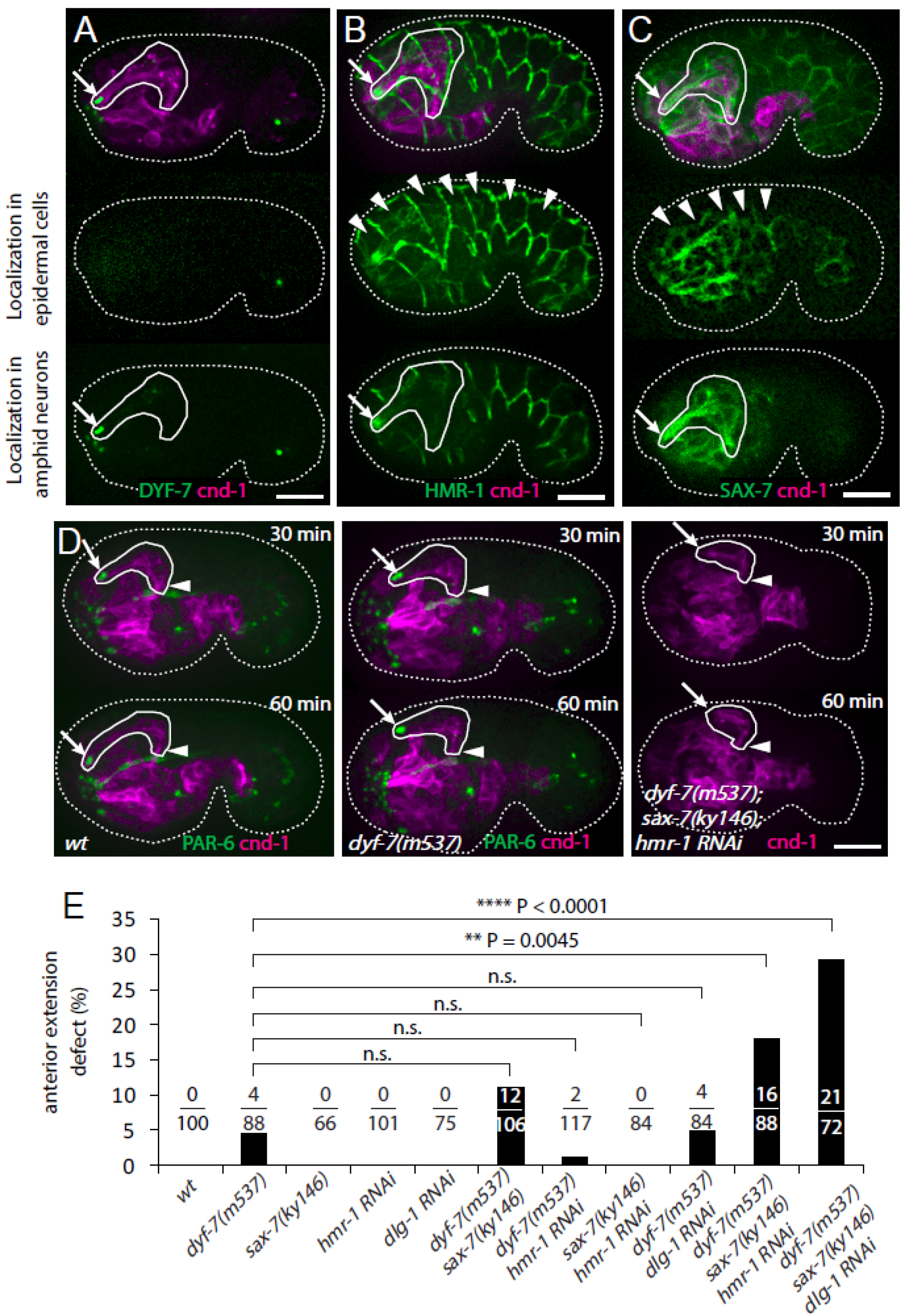
Multiple adhesion molecules function redundantly in amphid dendrite anterior extension. (A-C) Localization of DYF-7, HMR-1 and SAX-7 in amphid neurons and epidermal cells. Middle panels, superficial focal plane showing localization in epidermal cells. Lower panels, deeper focal plane showing localization in amphid neurons. Arrows indicate amphid tips and arrowheads indicate epidermal cells. (D) Dynamics of dendrite anterior extension in *wt, dyf-7(m537)* and *dyf-7;sax-7;hmr-1* triple loss of function. (E) Frequency of anterior extension defects in single, double and triple loss of function embryos. Number of amphids scored is indicated. P-values were calculated with Fisher’s exact test (two-tailed). Scale bars in A-D, 10 μm.

We then asked if these molecules are required for the attachment of the dendrite tips to the epidermis. If a gene is required, its loss of function will cause failure of attachment and result in partial or no anterior extension of the dendrites. In the majority of the *dyf-7(m537)* mutant embryos (84/88), the dendrite tip reached the sensory depression, but detached afterwards when the cell bodies moved posteriorly (Fig. S1), consistent with the known function of *dyf-7* in retrograde extension. However, in a small fraction of *dyf-7(m537)* embryos (4/88), the dendrites extended partially and the tip failed to reach the sensory depression (Fig. 3D). This result suggested that DYF-7 also plays a role in the attachment between epidermal cells and dendrite tips during the initial anterior extension. We did not find dendrite extension defects in *sax-7(kyl46), hmr-1(RNAi)* or *sax-7(ky146);hmr-1(RNAi)* embryos (Fig. 3E). Neither *sax-7(ky146)* nor *hmr-1(RNAi)* enhanced the phenotype of *dyf-7(m537)* in the anterior extension of the dendrites. However, in *dyf-7(m537);sax-7(ky146);hmr-1(RNAi)* triple loss of function embryos, we found a significant increase in anterior dendrite extension defects compared to *dyf-7(m537)* (Fig. 3E). This enhancement suggested that DYF-7 acts redundantly with SAX-7 and HMR-1 in the initial attachment of dendrite tips to the epidermis, before mediating the subsequent anchoring of dendrite endings at the embryonic nose as previously described. Similar results suggest that DLG-1 also functions in this process (Fig. 3E). In the epidermis, DLG-1 is known to regulate the recruitment and localization of HMR-1 to the junction (Firestein and Rongo, 2001; Koppen, et al., 2001).

### PAR-6 functions in amphid dendrite extension

To examine the function of PAR-6 in amphid dendrite extension, we generated *par-6(M/Z)* embryos that are deprived of both maternal and zygotic PAR-6 (Totong, et al., 2007). In this approach (see Methods), only a quarter of the embryos imaged in our experiments were expected to be *par-6(M/Z)*. We imaged a total of 89 embryos and found 25 to be *par-6(M/Z)* based on embryonic lethality. A previous study showed that PAR-6 is required for apical junction formation in epidermal cells by regulating the localization of HMR-1 and DLG-1 (Totong, et al., 2007). 17 of the 25 embryos arrested early (at 1.5-fold or soon after), indicating early malformation of the epidermis. Among these, 5 embryos showed partial dendrite extension (Fig. S2). This phenotype is likely due to the failure of the epidermis to complete its anterior migration. 8 of the 25 embryos developed to the 2-fold stage, suggesting that in these embryos epidermal morphogenesis was more or less normal beyond the stage where the dendrite tips normally reach the sensory depression. Among these 8 embryos, we found one in which the dendrites showed little extension (Fig. 4A). This result suggests that PAR-6 is required for the attachment of dendrite tips to the epidermis.

**Figure 4.**
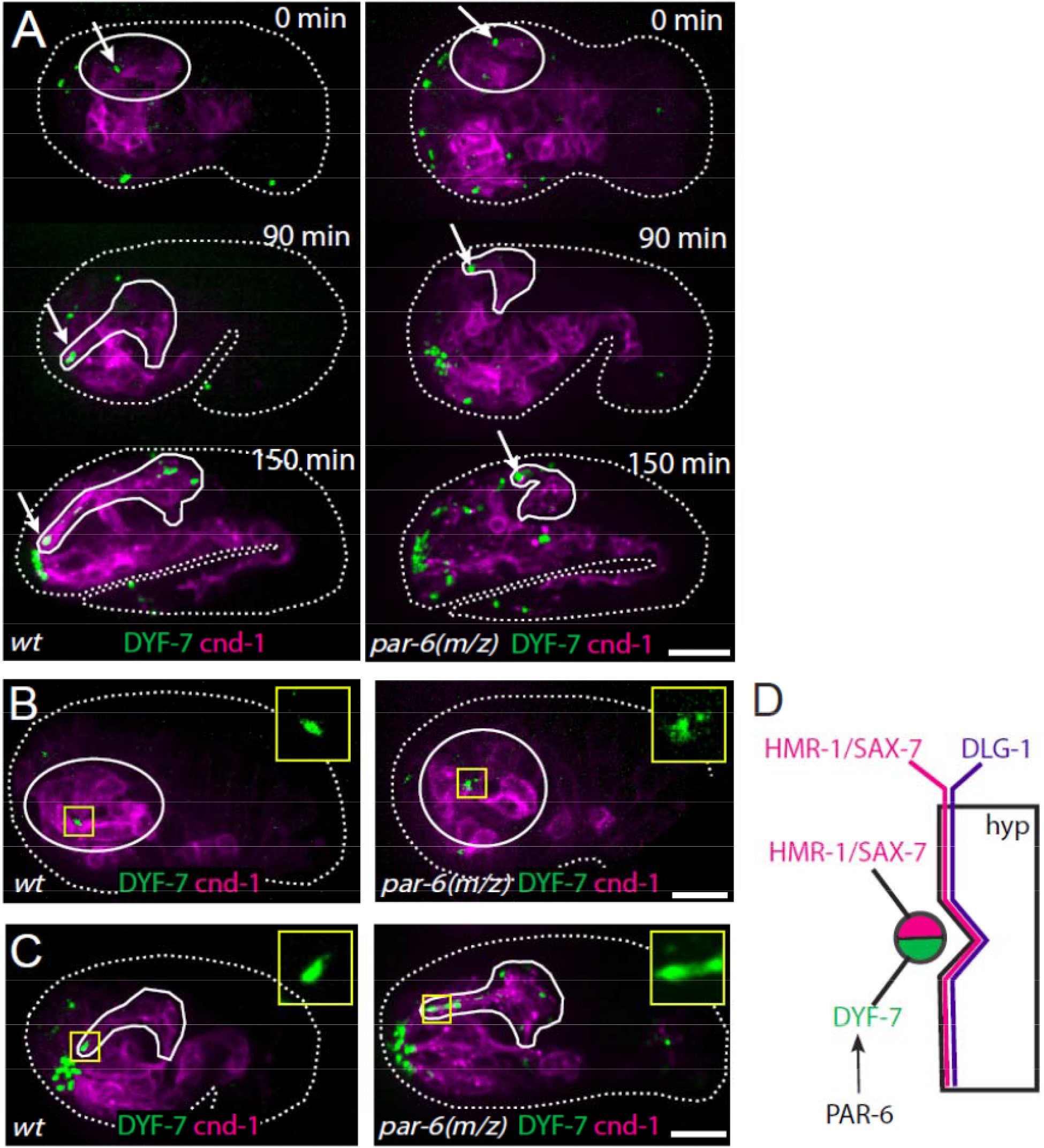
PAR-6 is required for amphid dendrite extension and regulates localization of DYF-7. (A) Abolished dendrite extension in *par-6(M/Z)* embryo. Arrows mark the dendrite tips. (B,C) DYF-7 shows more dispersed localization in *par-6(M/Z)*. (D) Schematic model of molecular localization between the amphid dendrite tips and epidermal cell. Scale bars in A-C, 10 μm.

Given the similar localization pattern and dendrite attachment phenotype of PAR-6 and DYF-7, we asked whether PAR-6 functions upstream of DYF-7. In the wild type, DYF-7 was condensed at the vertex of amphid rosette (Fig. 4B). The size of the DYF-7::GFP signal is 1.36 +/- 0.25 μm (mean +/- standard deviation, n=28). In 20% of the *par-6(M/Z)* embryos (5/25), DYF-7 showed a more diffused localization around the rosette center with the size of DYF-7::GFP signal greater than 3 standard deviations away from the wild-type mean (Fig. 4B). Furthermore, during dendrite extension, DYF-7 signal was spread along the dendrites in *par-6(M/Z)* embryos (7/25), instead of accumulating at the tip as in the wild type (Fig. 4C). These results suggested that PAR-6 regulates the localization of DYF-7 (Fig. 4D).

## DISCUSSION

Our results revealed that a multicellular rosette plays a central role in orchestrating multiple aspects of organogenesis of the *C. elegans* amphid. The collective extension of the dendrites and formation of the sensory nerve are mediated by the physical engagement of the sensory neurons at the rosette vertex, which demonstrate a mechanism of collective neurite extension that is different from the classical pioneer-follower model. Furthermore, the rosette vertex is coupled to the migrating epidermis and carried anteriorly to extrude the dendrites, which provides a mechanism to coordinate dendrite growth with the morphogenesis of surrounding tissues that participate in head formation. More broadly, key structural features of the mature amphid are already seeded in the rosette, which includes polarization of neurons and future dendrite tips at the vertex, engagement of the future dendrite tips with the amphid glia cells which ultimately ensheath the dendrites, and engagement of the future dendrite tips with the epidermis. Thus, topological features of an organ can be specified early via short range interactions when players are local, before these features are extended over space through development.

Multicellular rosettes have been found in diverse developmental contexts to orchestrate tissue morphogenesis (Harding, et al., 2014). In *C. elegans*, we recently showed that dynamic formation and resolution of multicellular rosettes mediate cell intercalation and convergent extension during assembly of the ventral nerve cord (VNC) (Shah, et al., 2017). It is not clear what makes the amphid rosette stable while those in the VNC are dynamic. One possibility is the different cell polarity pathways used in the rosettes, which may regulate cell adhesion differently. In the amphid rosette, PAR-6 is localized to the vertex. In the VNC rosettes, VANG-1/Van Gogh and SAX-3/Robo are localized to the vertex and are required for timely rosette resolution. However, two lines of evidence argue against this simple model. First, in *Drosophila* germband extension where dynamic rosette-based cell intercalation was first characterized (Blankenship, et al., 2006), PAR-3 localizes to non-contracting cell-cell contacts (Pare, et al., 2014), suggesting that the PAR polarity pathway does not necessarily lead to stable rosettes. Second, in our study, *par-6(M/Z)* did not cause rosette disassembly. Similarly, the use of DYF-7 in the amphid rosette also does not offer a simple explanation of its stability. Loss of function of *dyf-7*, either alone or in combination with *sax-7* and *hmr-1* did not cause rosette disassembly.

Surrounding tissues play profound roles in neuronal development (Chao, et al., 2009), and recent work has highlighted intriguing roles for skin in shaping dendrites. In *Drosophila*, signals from the skin shape Class IV da neurons (Meltzer, et al., 2016; Jiang, et al., 2014; Parrish, et al., 2007), and skin acts to clear debris from degenerating neurons in zebrafish (Rasmussen, et al., 2015). In *C. elegans*, the epidermis plays a role in controlling synapse density (Cherra and Jin, 2016) and positioning (Shao, et al., 2013). Furthermore, the epidermis plays a critical role in patterning the elaborate dendritic arbor of the PVD neuron. Interestingly, SAX-7/L1CAM functions in this interaction. Specifically, it functions together with MNR-1 in the epidermis to instruct the dendritic pattern of PVD (Dong, et al., 2013; Salzberg, et al., 2013). In our study, it remains unclear whether SAX-7 acts in the neurons, skin, or both, but the interaction between the amphid dendrites and the epidermis appears to be a simple physical attachment at a focal point defined by the rosette. This example therefore suggests that skin not only provides signaling cues to developing neurons, but can also provide mechanical cues that shape dendrites by physically coupling neurons to epidermal morphogenesis.

We showed that multiple adhesion molecules function redundantly with DYF-7 to mediate the attachment of the rosette vertex and dendrite tips to the epidermis. The DLG-1 reporter used in our study showed expression only in the epidermis and a previous study by immunostaining suggested that DLG-1 is absent in neurons (Firestein and Rongo, 2001). However, a more recent study showed that AJM-1, which functions with DLG-1 in the apical junction, is localized at the dendrite tips of amphid neurons in postembryonic worms (Nechipurenko, et al., 2016). Further investigation is required to determine if DLG-1 is present at the dendrite tips during extension. Recent work also suggested that transition zone proteins in cilia promote dendrite development in related sensory neurons in the tail of the worm (Schouteden, et al., 2015). However, transition zone proteins are not present at the amphid dendrite tips until the 2-fold stage (Schouteden, et al., 2015), about an hour after the rosette-mediated anterior extension is complete.

PAR-6 localization to the dendrite tips raises an interesting contrast to its localization and function in vertebrate neurons. In cultured hippocampal neurons, the PAR-6/PAR-3/aPKC complex is enriched at the tip of the future axon. The complex is thought to provide local regulation of the actin cytoskeleton (Insolera, et al., 2011). A genetic screen in *C. elegans* for mutants that disrupt the axon-dendrite polarity identified *unc-33*, which encodes a microtubule binding protein called CRMP (Maniar, et al., 2011). UNC-33/CRMP localizes to the initial segment of the axon and has global effects on microtubule organization in a neuron. The potential function of PAR-6 in the axon-dendrite polarity in the amphid neurons has not been examined. Intriguingly, in *Drosophila* sensory and motor neurons, PAR-6/PAR-3/aPKC localizes to dendrites (Sanchez-Soriano, et al., 2005). It remains to be seen if this similarity is a coincidence or marks a difference between vertebrate and invertebrate neurons.

## EXPERIMENTAL PROCEDURES

### Worm Stains and Genetics

Worms were maintained at room temperature on nematode growth media (NGM) plates seeded with 0P50 bacteria as previously described (Brenner, 1974), except for RNAi experiments. N2 Bristol was used as the wild-type strain. All strains are listed in Table S1.

### RNAi

RNAi experiment was performed using the standard feeding method (Rual, et al., 2004; Kamath, et al., 2003). For *elt-1(RNAi)* and *hmr-1(RNAi)*, L1 hermaphrodites were placed on RNAi plates and embryos were harvested by cutting the adults 2 days later. For *dlg-1(RNAi)*, L4 hermaphrodites were grown for 1 day before embryos are collected. The clones of 10019-B-8 from Vidal RNAi library, I-5F23 and X-8A08 from Ahringer RNAi library were used to target *elt-1, hmr-1* and *dlg-1*, respectively.

### Microscopy

Embryos were collected and mounted as previously described (Bao and Murray, 2011). Briefly, embryos were collected by cutting gravid hermaphrodites in a droplet (20 μL) of M9 buffer (3 g KH_2_PO_4_, 6 g Na_2_HPO_4_, 5 g NaCl, 1 ml 1 M MgSO_4_, per liter H_2_O). Embryos at the 2 or 4-cell stages were transferred to a droplet (1.5 μL) of M9 containing 20 μm polystyrene beads on 24 × 50 mm coverglass. An 18 × 18 mm coverglass was laid on top and sealed using melted Vaseline. Images were acquired on a spinning disk confocal microscope (Quorum technologies) comprising a Zeiss Axio Observer Z1 frame. An Olympus UPLSAPO 60xs objective was used with a thread adapter to mount on the Zeiss body (Thorlabs). The timing of major developmental events was used to check for phototoxicity during 3D time-lapse imaging.

## AUTHOR CONTRIBUTIONS

LF, IK and ZB designed the experiments. LF and IK performed the experiments and MGH constructed reagents. LF, IK and ZB interpreted the results. LF drafted the manuscript, and ZB and MGH edited it. All authors reviewed the manuscript.

## ACKNOWLEDGMENTS

The authors would like to thank S. Shaham for reagents and discussions, D. Colon-Ramos and Songhai Shi for advice on the manuscript, and members of the Bao lab for discussions and help. This work was supported by NIH grants (R01 GM097576 and R24 OD016474 to ZB) and the MSK Cancer Center Support/Core (P30 CA008748). Some strains were provided by Dr. Jeremy Nance, Dr. Shohei Mitani (National Bioresource Project), and the CGC, which is funded by the NIH Office of Research Infrastructure Programs (P40 OD010440).

## SUPPLEMENTAL INFORMATION

Supplemental Information includes one table, two figures and one movie.

**Table S1.** *C. elegans* strains used in this study, related to Experimental Procedures

|  |  |
| --- | --- |
| BV293 | zbIs3[cnd-1p::PH::GFP]; ujIs113[pie-1p::mCherry::H2B; nhr-2p::mCherry::HIS-24-let-858UTR; unc-119(+)]II. |
| JIM113 | ujIs113[pie-1p::mCherry::H2B; nhr-2p::mCherry::HIS-24-let-858UTR; unc-119(+)]II. |
| BV294 | itIs1024[par-6::PAR-6::GFP]; zyIs36[cnd-1p::PH::mCherry; myo-2p::mCherry]X. |
| KK1024 | itIs1024[par-6::PAR-6::GFP] |
| OU412 | zyIs36[cnd-1p::PH::mCherry; myo-2p::mCherry]X |
| BV300 | dyf-7 (ns119) X; hmnEx149[dyf-7p::DYF-7(ZP-sfGFP)-mCherry; rol-6(su1006)]; zyIs36[cnd-1p::PH::mCherry; myo-2p::mCherry]X. |
| CHB440 | dyf-7 (ns119) X; hmnEx149[dyf-7p::DYF-7(ZP-sfGFP)-mCherry; rol-6(su1006)] |
| BV403 | mcIs40 [lin-26p::ABDvab-10::mCherry; myo-2p::GFP]; zyIs36[cnd-1p::PH::mCherry; myo-2p::mCherry]X. |
| ML916 | mcIs40 [lin-26p::ABDvab-10::mCherry; myo-2p::GFP] |
| BV496 | xnIs17[dlg-1::GFP; rol-6(su1006)]; axIs1928[mCherry::PAR-6] |
| FT63 | xnIs17[dlg-1::GFP; rol-6(su1006)] |
| JH2647 | axIs1928 [mCherry::PAR-6]; axIs1182[GFP::PAR-2]. |
| BV337 | unc-119(ed3) III; ddIs290[sax-7::TY1::EGFP::3xFLAG(92C12); unc-119(+)]; zyIs36[cnd-1p::PH::mCherry; myo-2p::mCherry]X. |
| TH502 | unc-119(ed3) III; ddIs290[sax-7::TY1::EGFP::3xFLAG(92C12); unc-119(+)]. |
| BV308 | unc-119(ed3) III; xnIs96 [hmr-1p::HMR-1::GFP::unc-54 3'UTR; unc-119(+)]; zyIs36[cnd-1p::PH::mCherry; myo-2p::mCherry]X. |
| FT250 | unc-119(ed3) III; xnIs96 [hmr-1p::HMR-1::GFP::unc-54 3'UTR; unc-119(+)]. |
| BV363 | dyf-7(m537) zyIs36[cnd-1p::PH::mCherry; myo-2p::mCherry]X; ntIs1[gcy-5::GFP]; itIs1024[par-6::PAR-6::GFP]. |
| BV341 | dyf-7(m537) zyIs36[cnd-1p::PH::mCherry; myo-2p::mCherry]X; ntIs1[gcy-5::GFP]. |
| OS611 | dyf-7(m537)X; nsIs53[vap-1::RFP]IV; ntIs1[gcy-5::GFP]. |
| BV392 | dyf-7(m537) zyIs36[cnd-1p::PH::mCherry; myo-2p::mCherry]X; ntIs1[gcy-5::GFP]; sax-7(ky146)IV. |
| CX2993 | sax-7(ky146) IV; kyIs4[ceh-23-unc-76-gfp::lin-15]X. |
| BV425 | sax-7(ky146) IV; zyIs36[cnd-1p::PH::mCherry; myo-2p::mCherry]X. |
| BV309 | par-6(tm1425)/unc-101(m1)I; him-8(e1489)IV; zyIs36[cnd-1p::PH::mCherry; myo-2p::mCherry]X. |
| JJ1743 | par-6(tm1425)/hIn1[unc-54(h1040)]I; him-8(e1489)IV. |
| BV295 | unc-101(m1)I; zyIs36[cnd-1p::PH::mCherry; myo-2p::mCherry]X. |
| DR1 | unc-101(m1)I. |
| BV307 | unc-101(m1) par-6(zu170)I; zuIs43[pie-1::GFP::PAR-6::ZF1; unc-119(+)]; zyIs36[cnd-1p::PH::mCherry; myo-2p::mCherry]X. |
| FT36 | unc-101(m1) par-6(zu170)I; zuIs43[pie-1::GFP::PAR-6::ZF1; unc-119(+)] |
| BV420 | unc-101(m1) par-6(zu170)I; zuIs43[pie-1::GFP::PAR-6::ZF1; unc-119(+)]; zyIs36[cnd-1p::PH::mCherry; myo-2p::mCherry]X; hmnEx149[dyf-7p::DYF-7(ZP-sfGFP)-mCherry; rol-6(su1006)]. |

**Figure S1.**
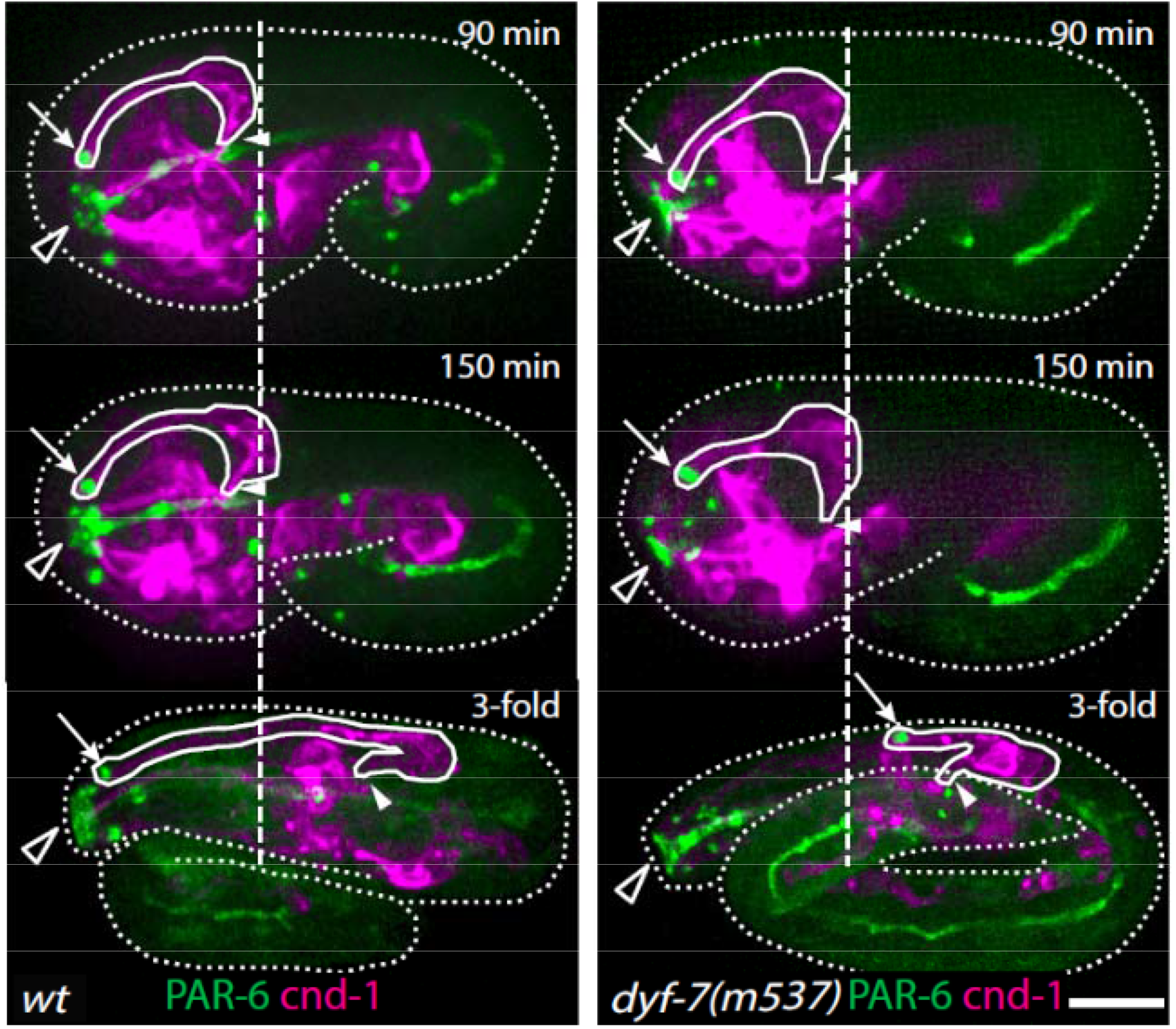
Retrograde extension and defects in *dyf-7(m537)*. Left: In the wild type, after the initial anterior extension is complete and the dendrite tips reach the sensory depression (top panel), cell bodies start to move posteriorly as the embryo elongates, further extending the dendrites through retrograde extension. Right: In most of *dyf-7(m537)* embryos, the dendrite tips detach from the sensory depression during embryo elongation, after successful anterior extension. Arrows indicate amphid dendrite tips. Closed arrowheads indicate amphid commissure. Open arrowheads indicate the sensory depression. Dashed line mark the posterior end of the amphid neurons at the time when anterior dendrite extension is complete. Scale bar, 10 μm.

**Figure S2.**
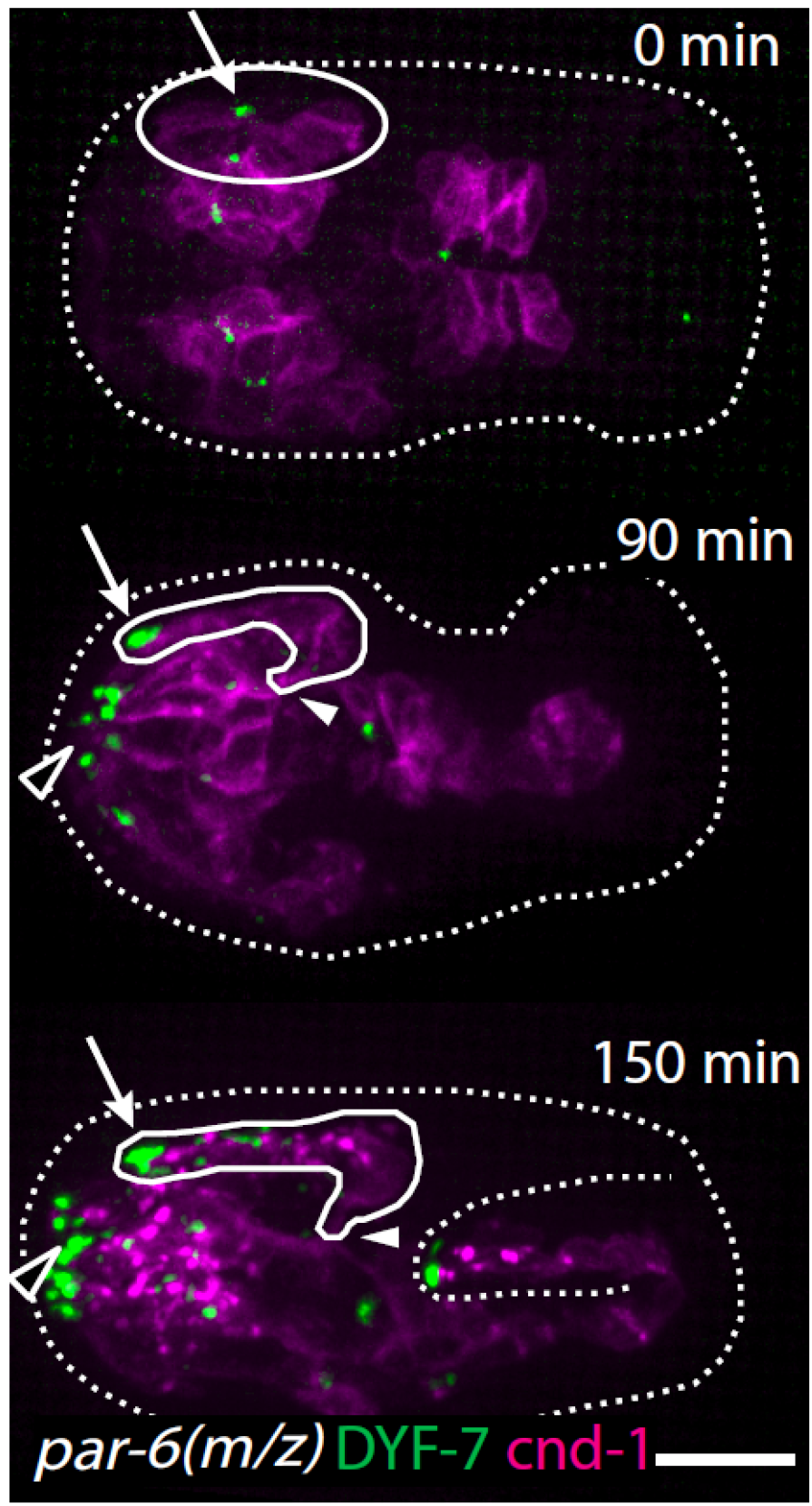
Partial extension of amphid dendrites in *par-6(M/Z)* embryos that arrested at the 1.5-fold stage. See Figure 4 for the wild-type control. Arrows indicate amphid dendrite tips. Closed arrowheads indicate amphid commissure. Open arrowheads indicate the sensory depression. Scale bar, 10 μm.

Movie S1. Amphid dendrite anterior extension. Time-lapse imaging of a wild-type embryo expressing *cnd-1*p::PH::mCherry to label sensory neurons and PAR-6::GFP to label dendrite tips. Dashed lines mark the initial position of the amphid neuron cell bodies at 0 min. The dendrite tip is tracked with an arrow. Scale bar, 10 μm.

